# Biodiversity effects of beaver activity in a semi-natural enclosure revealed by eDNA

**DOI:** 10.64898/2026.05.15.725411

**Authors:** B Hänfling, NP Griffiths, JA Macarthur, B Morrissey, D Svobodova, V Pritchard, A Tree, M Gaywood

## Abstract

1. Environmental DNA (eDNA) metabarcoding is an emerging tool for biodiversity assessment in freshwater systems, offering high-resolution insights into community composition. Here, we apply eDNA metabarcoding to evaluate the ecological impacts of Eurasian beaver (*Castor fiber*) activity within a seminatural enclosure in the Scottish Highlands.
2. We collected seasonal water samples from nine sites, six influenced by beaver dams and three control sites with no evidence of beaver engineering, across a 40-hectare enclosure. Samples were analysed for vertebrate and macroinvertebrate diversity using established 12S and COI markers.
3. Vertebrate alpha diversity did not differ significantly between beaver and control sites, likely reflecting the small spatial scale and low species richness of upland Scottish streams. However, community composition differed significantly between treatments, especially for fish (PERMANOVA, R^2^ = 0.55, *P* < 0.001), with beaver-influenced sites dominated by three-spined stickleback and control sites by brown trout. Macroinvertebrate communities showed a 78% increase in gamma diversity in beaver-modified habitats relative to controls. Species composition varied strongly with beaver presence (PERMANOVA, R^2^ = 0.29, *P* < 0.001), likely due to the creation of lentic-lotic mosaics and associated microhabitat diversity. Seasonal variation was significant in both taxonomic groups, with the lowest species richness and highest community dispersion observed in summer, probably reflecting hydrological and temperature-driven dynamics in eDNA production and transport.
4. Our findings reinforce previous evidence that beaver dam-building activity enhances beta diversity in headwater systems. Additionally, we demonstrate that eDNA metabarcoding is a sensitive method for detecting spatial patterns in freshwater biodiversity associated with these activities at scales ranging from tens to hundreds of meters. These approaches could inform future monitoring strategies aligned with landscape-scale beaver management and reintroductions.

## Introduction

The Eurasian beaver (*Castor fiber*) population experienced a significant decline during the early last millennium but has since shown notable recovery due to conservation translocation efforts and natural recolonisation across Europe (Campbell-Palmer *et* al., 2016; Halley, Saveljev & Rosell, 2021). The North American beaver (*Castor canadensis*) is also expanding its range due to changing environmental conditions (Tape et al., 2018). Interest is also growing in the restoration of the Eurasian beaver within Britain. Beavers, regarded as ecosystem engineers, transform their environments by felling trees, constructing dams, creating nutrient-rich ponds, and slowing water flow (Campbell-Palmer et al., 2016; Law et al., 2017). This process results in the creation of complex habitats, a key motivation behind their reintroduction, as it has demonstrated positive effects on biodiversity in various studies (Wright, Jones & Flecker, 2002; Kemp et al., 2012; Gaywood et al., 2015; Law, McLean & Willby, 2016; Stringer & Gaywood, 2016). However, the full implications for biodiversity and ecosystem services in a British context are not yet fully understood and there are multiple potential conflicts with other forms of land use such as agriculture, forestry, and recreational fishing (Stringer, Blake & Gaywood, 2015). Beaver activity is highly dynamic, and impacts are therefore predicted to be dependent on the surrounding landscape, and to change over time. There have been some case studies that have looked at biodiversity impacts of beaver in Scottish freshwaters and those have largely confirmed that beaver wetland creation has a positive impact on the diversity of plants and aquatic invertebrates (Law et al., 2016, 2017; Needham et al., 2021). The Scottish beaver trial at Knapdale observed increasing diversity of standing water plant and water beetle species, although specific plant species were negatively affected (Willby, Perfect & Law, 2014). No effect on otter presence was recorded during the Scottish Beaver Trial (Harrington et al., 2015). Preliminary studies on fish presence during the Scottish Beaver Trial did not record changes in the fish community (Argyll Fisheries Trust, 2015), and a separate study at a Scottish Highlands enclosed site found beavers had profound effects on the local brown trout (*Salmo trutta*) that promoted higher abundances of larger size classes (Needham et al. 2021). However, to date these Scottish studies have been constrained to loch systems and small tributaries in the headwater region. A more holistic understanding of the effects of beaver on biodiversity both in terms of habitats and taxonomic groups studied, is needed in order to determine where beaver reintroductions are likely to be most beneficial, or potentially detrimental to biodiversity (IUCN/CPSG, 2022) or where the cost-benefit trade-off justifies population regulation of a natural population. This requires efficient and effective biodiversity monitoring strategies which can be upscaled to a catchment level.

Environmental DNA (eDNA) metabarcoding approaches are increasingly used for biodiversity monitoring of aquatic habitats. This approach combines eDNA with modern High-Throughput-Sequencing technology and allows the simultaneous characterization of entire biological communities. Previous research has shown that eDNA metabarcoding is more effective at detecting elusive fish species than established invasive surveying techniques such as electric fishing or fyke netting, and in both lentic and lotic habitats (Hänfling et al., 2016; Pont et al., 2018; Griffiths et al., 2020; Czeglédi et al., 2021). eDNA metabarcoding approaches have also been successfully used to characterise aquatic invertebrate communities (Elbrecht & Leese, 2015; Blackman et al., 2019; Harper et al., 2021) and to detect cryptic non-native invertebrate species (Blackman et al., 2017). However, eDNA metabarcoding results for invertebrates are usually not directly comparable to conventional survey methods such as kick-sampling as the taxonomic overlap between the different approaches is limited (Gleason et al., 2021; Harper et al., 2021). Correspondingly, the use of these data for ecological status monitoring requires a re-evaluation of the quality indices used (Hering et al., 2018). For fish, eDNA metabarcoding can also provide reliable semi-quantitative estimates and in lentic habitats, the approach is now being deployed in routine monitoring programmes by the UK government agencies and to estimate the ecological status of lakes (Willby et al., 2019).

eDNA metabarcoding offers a promising method for generating the biodiversity data needed to inform management strategies for expanding beaver populations in both Eurasia and North America. For example, in anticipation of further conservation translocations and the expansion of the Scottish beaver population into new catchments, the Scottish Beaver Strategy 2022–2027 was developed (IUCN/CPSG, 2022). A key goal of this strategy is the ongoing assessment of the implications of beaver presence impacts on biodiversity, with a specific action to develop technologies that can assess interactions with priority terrestrial and aquatic species and communities and other species. This project set out to evaluate whether environmental DNA (eDNA)-based methods are suitable for supporting such monitoring objectives.

The objectives of this study were to (i) characterise vertebrate and invertebrate communities at beaver-affected sites within a semi-natural beaver enclosure across different seasons, using eDNA metabarcoding of water samples, and (ii) compare biodiversity patterns between sites influenced by beaver activity and control sites with no visible beaver engineering. Based on prior conventional surveys in this enclosure, we hypothesised that (i) trout eDNA concentrations would be lower in the beaver-influenced wetland compared to control areas, (ii) total macroinvertebrate species richness would be higher in beaver-modified areas, and (iii) the presence of beaver wetlands would enhance beta diversity across the study site.

Addressing these objectives will also provide insights into monitoring the effectiveness of other conservation and restoration projects.

## Methods

### Experimental design

The study was conducted in a semi-natural beaver enclosure located in the Scottish Highlands, within the headwaters of the Moniack Burn catchment. Established in 2008, the 40-hectare enclosure has supported a small population of 2–5 beavers since its inception. The site features a small artificial loch fed by two streams of roughly equal size and character, as well as a single outflow stream. One of the inflow streams (hereafter “beavers above loch”) and the outflow stream (hereafter “beavers below loch”) have been extensively modified by beaver activity since 2009 and 2016, respectively, forming a lentic-lotic mosaic (Figures 1, 2). These modified streams now contain multiple beaver dams and expansive wetland areas that support a diversity of aquatic and terrestrial habitats. The second inflow stream, which has not been physically altered by beavers, retains its original lotic character and serves as a control (hereafter “control stream”). Water samples were collected from nine sites across the study area, three within each of the three spatial groupings (beavers above loch, control stream, and beavers below loch; see Figure 1). In 2022, 81 2-litre surface water samples were collected across 9 sites (3 replicates per site) following the methodology described in Griffiths *et al*. (2025) (Figure 1). Sampling was conducted on three occasions in 2022 to capture seasonal variation: 28–29 March (spring), 20–21 June (summer), and 24–25 October (autumn).

**Figure 1:**
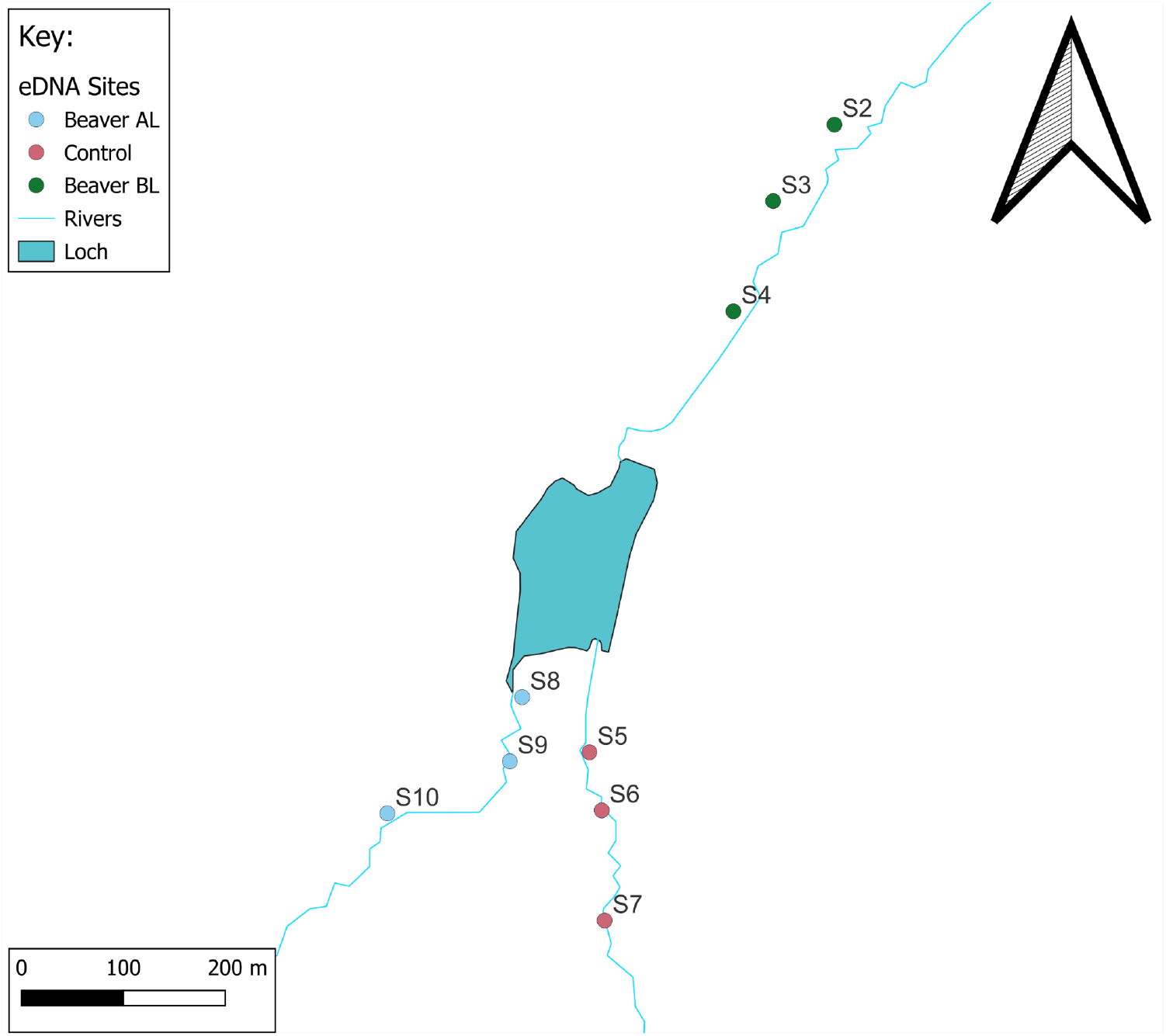
Schematic representation of the eDNA sample sites at the enclosure, sites are coloured by their group (Beavers above loch = blue, Beavers below loch = green, Control sites = red).

### Water sample collection

At each site, three 2 litre replicate water samples were collected and filtered within 24 hours at the eDNA laboratory at UHI Inverness following the methodology described in Griffiths *et al*. (2025). Standard Operating Procedures for water collection include strict protocols for avoiding contamination (e.g. bleach treatment of equipment, gloves etc.) and detecting potential contamination (control samples and blanks at all stages of the workflow). See Annex 1 for a detailed methodology.

**Figure 2:**
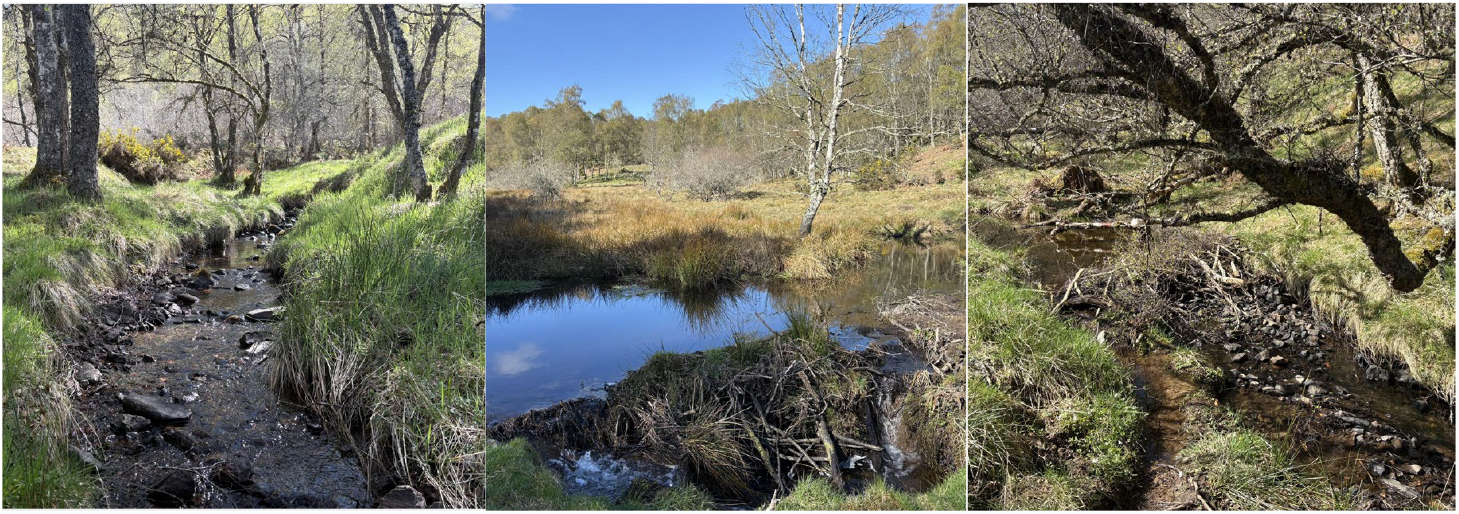
Typical habitats in the beaver enclosure. Lotic habitat in the control stream (left); lentic and lentic lotic habitat in the beaver modified sections (middle and right).

### Molecular methods

Established eDNA metabarcoding workflows were used to analyse vertebrate and invertebrate eDNA (See Annex 1 for full methodological details). Vertebrate metabarcoding was carried out using a vertebrate specific 12S marker (Riaz *et al*., 2011), which has previously been shown to produce reproducible, semi-quantitative data for UK freshwater fish (Hänfling *et al*., 2016) and to amplify all amphibian, mammal, and bird DNA reliably, but not reptiles. The taxonomic resolution of the 12S marker is largely to species level for fish, mammals and amphibians with a few exceptions where only genus level assignments are possible. The taxonomic resolution is very variable for birds with several key groups only identifiable to family level, such as gulls (Laridae), ducks (Anatidae) and crows (Corvidae). Bird data were therefore not analysed in detail. Invertebrate metabarcoding was carried out using a molecular marker based on the mitochondrial cytochrome oxidase I gene (COI), which was designed to work well for key arthropod groups traditionally used in freshwater monitoring such as mayflies, chironomids and beetles as well as other key invertebrate groups (Leese *et al*., 2021). There is currently no evidence that this approach can be used to assess the relative abundance of invertebrate species.

### Bioinformatics and data analysis

Raw sequence data were analysed using the bioinformatics pipeline Tapirs (https://github.com/EvoHull/Tapirs). After quality control, query sequences were then compared against the reference database using *BLAST (Zhang et al., 2000)*. Taxonomic identity was assigned using a custom majority lowest common ancestor (MLCA) approach based on the top 2% query *BLAST* hit bit-scores, with at least 90% query coverage and a minimum identity of 95% for invertebrates and 98% for vertebrates. All statistical analyses and data visualisation was carried out using the statistical programming environment R v.4.5.2 (R Core Team, 2025). Full methodological details are provided in Annex 1.

Following the bioinformatics, a minimum threshold of five reads was applied to the vertebrate dataset to remove low-frequency reads alongside a species-specific contaminant threshold to remove any reads in samples which were lower than detections in controls (Figure S1) (Macarthur et al., 2025). The invertebrate dataset had a low-frequency reads threshold of <20 reads applied, to reduce sporadic detections (Griffiths et al., 2024). Subsequently, field replicates were pooled prior to downstream analyses.

For fish and amphibians, the number of sequence reads assigned to species were converted to relative reads per sample (proportional reads) to create a standardised measure of eDNA community composition across samples (species reads / total sample reads). Previous research has found a strong correlation of fish read counts with actual recorded abundance and biomass of fish communities within UK freshwater systems (Li et al., 2019; Di Muri et al., 2022); and these data can therefore be interpreted as a proxy of relative abundance. Therefore, the compositional dissimilarity of fish and amphibian communities of individual sites was analysed using the Bray-Curtis dissimilarity index. Mammal abundance can be more difficult to quantify from eDNA, due to the sporadic nature of their interactions with water. Similarly, no such relationship has been demonstrated for invertebrate metabarcoding data. Therefore, for mammals and invertebrates the Jaccard index was chosen as a conservative approach which takes presence/absence to indicate the occurrence of a species within a water sample.

Alpha diversity was calculated as the average number of species per water sample (species richness). To assess spatial and seasonal variation in alpha diversity, nonparametric Kruskal-Wallis tests from the package stats v3.6.3 were used to compare alpha diversity of communities between groups and seasons (Harper et al., 2020). The total number of species for each of the three spatial groupings across all seasons was used as a measure for Ɣ – diversity.

The DECOSTAND function in the package vegan v2.5-7 (Oksanen et al., 2020) was used to convert the read counts into presence/absence or proportional read counts for statistical analyses of beta diversity. To assess variation in community composition across the groupings, Jaccard and Bray-Curtis dissimilarity matrices were created using the VEGDIST function and visualised via Non-Metric Multidimensional Scaling (NMDS) using the METAMDS function (Harper *et al*., 2020). A series of PERMANOVAs were then carried out to statistically test for spatial and temporal differences in the community composition for each group. An interaction term was included in each PERMANOVA to capture the combined effect of spatial and temporal variation.

Alongside all PERMANOVAs, the homogeneity of multivariate dispersions (MVDISP) was calculated using the BETADISPER function in vegan v2.5-7 and statistically tested using ANOVA. This was used to verify whether a significant result was due to a difference in the mean community composition as opposed to a difference in dispersion between samples. All PERMANOVA were performed with a Jaccard or Bray-Curtis distance matrix and 999 permutations, using the function “adonis2” in the Vegan package (Oksanen et al., 2020).

## Results

### Vertebrate diversity

A total of 1.45m, 1.74m and 2.21m DNA sequences could be assigned to vertebrate taxa in the spring, summer and autumn data sets respectively. The proportion of sequences assigned to different vertebrate groups varied across seasons but was consistently highest for fish. Reads assigned to target species (other than human) in control samples, including field blanks, extraction blanks, PCR negatives, and PCR positives, were low. Across 33 control samples, only 7 reads of *G. aculeatus* and 42 reads of *S. trutta* were detected, each occurring in a single sample (Figure S1).

Across all seasons 25 vertebrate taxa were detected (excluding birds and reptiles), three amphibians, four fish species and 18 mammal species (Fig. 3). The total species richness was almost identical among groups with 21, 22 and 19 species detected across the beavers AL, beavers BL and control respectively. However, the overall number of occurrences (i.e. sum of all positive detections across all species not including beavers) was higher in the beaver sites with 168 and 160 occurrences for Beavers AL and Beavers BL respectively and 143 for the Control site. Most species were detected across all three spatial groups but with some notable exceptions. Beaver eDNA was not detected in the control stream confirming the absence of the species in this site. European Eels (*Anguilla anguilla*) were only detected in the beaver sites and not detected in any sample from the control. Water vole (*Arvicola amphibius*), water shrew (*Neomys fodiens*) and palmate newt (*Lissotriton helveticus*) eDNA were all detected in a higher proportion of beaver-modified sites compared to the control. Other species which were only detected in some of the spatial groups were rare species with only single occurrences. Additionally, some more common species varied in read counts and occurrences among groups.

**Figure 3:**
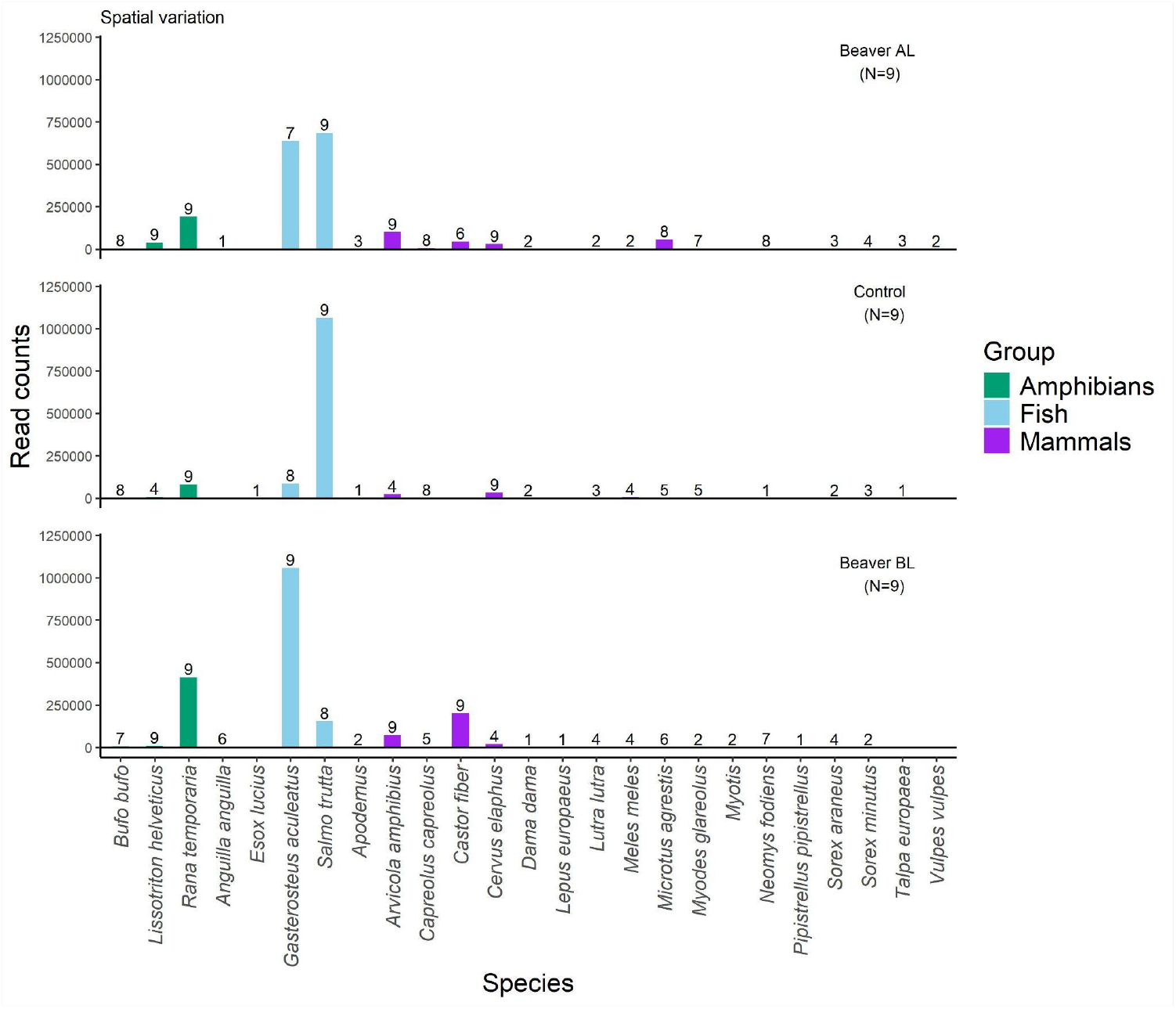
Number of read counts for individual vertebrate taxa in samples from the three different spatial groups; Beaver dammed sites above loch (top), control site above loch (middle) and beaver dammed sites below loch (bottom). 100% = 9 occurrences. Birds were omitted from this graph, due to the lack of taxonomic resolution in this group

There was also considerable variation in sequence read counts for individual species among the three spatial groups (Fig. 3). The control stream was largely dominated by brown trout eDNA, whereas the sites with beaver dams contained considerably more stickleback and amphibian eDNA. Water vole (*A. amphibius*) eDNA was also present in a higher proportion in sites with beaver dams compared to the control site.

Mean alpha diversity of samples did not differ significantly between the three spatial groups (Fig 4a; Kruskal-Wallis: X2 = 1.97, DF=2, P=0.374) but was significantly different between seasons with lowest species richness being recorded in summer (Fig 4b; Kruskal-Wallis: X2 = 12.998, DF=2, P=0.002).

**Figure 4:**
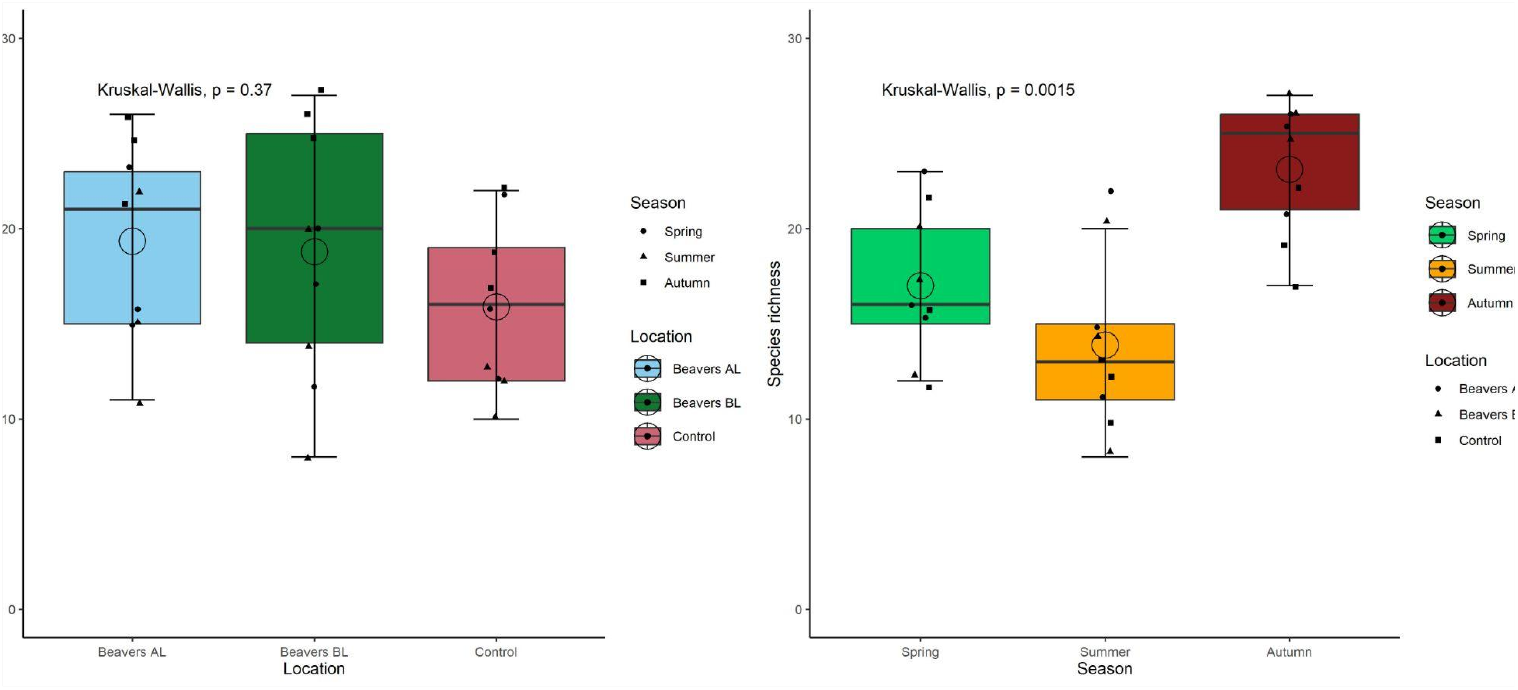
Box plots comparing total vertebrate species richness in samples among different spatial groups (left) and different sampling seasons (right).

Community composition varied significantly for all three taxonomic groups between beaver and control sites (Fig. 5, Table 1), but the effect was particularly strong for fish (R2=0.55, P=0.001) compared to mammals (R2=0.24, P=0.001) and Amphibians: R2=0.21, P=0.001). Multivariate dispersions were homogeneous between groups (Table 1).

**Table 1:** Analysis of community similarity among spatial groups and seasons for different vertebrate groups.

|  | Homogeneity of multivariate dispersions (ANOVA) |  |  | Community similarity (PERMANOVA) |  |  |  |
| --- | --- | --- | --- | --- | --- | --- | --- |
| Covariate | df | F | P | df | R2 | F | P |
| **Mammals** |  |  |  |  |  |  |  |
| Location | 2 | 3.03 | 0.067 | 2 | 0.24 | 4.61 | 0.001 *** |
| Season | 2 | 5.68 | 0.01 ** | 2 | 0.15 | 2.83 | 0.001 *** |
| Location * Season | n/a | n/a | n/a | 4 | 0.15 | 1.42 | 0.09 |
| **Fish** |  |  |  |  |  |  |  |
| Location | 2 | 2.79 | 0.081 | 2 | 0.55 | 13.70 | 0.001 *** |
| Season | 2 | 0.35 | 0.707 | 2 | 0.03 | 0.87 | 0.431 |
| Location * Season | n/a | n/a | n/a | 4 | 0.05 | 0.62 | 0.679 |
| **Amphibians** |  |  |  |  |  |  |  |
| Location | 2 | 0.21 | 0.811 | 2 | 0.21 | 4.77 | 0.001 *** |
| Season | 2 | 1.31 | 0.29 | 2 | 0.16 | 3.71 | 0.006 *** |
| Location * Season | n/a | n/a | n/a | 4 | 0.24 | 2.83 | 0.007 *** |

Community composition varied also significantly but moderately for mammals and amphibians between seasons but not for the fish (Fig. 5, Table 1). Multivariate dispersions were homogeneous between seasons for fish and amphibians, but not for mammals (Table 1).

**Figure 5:**
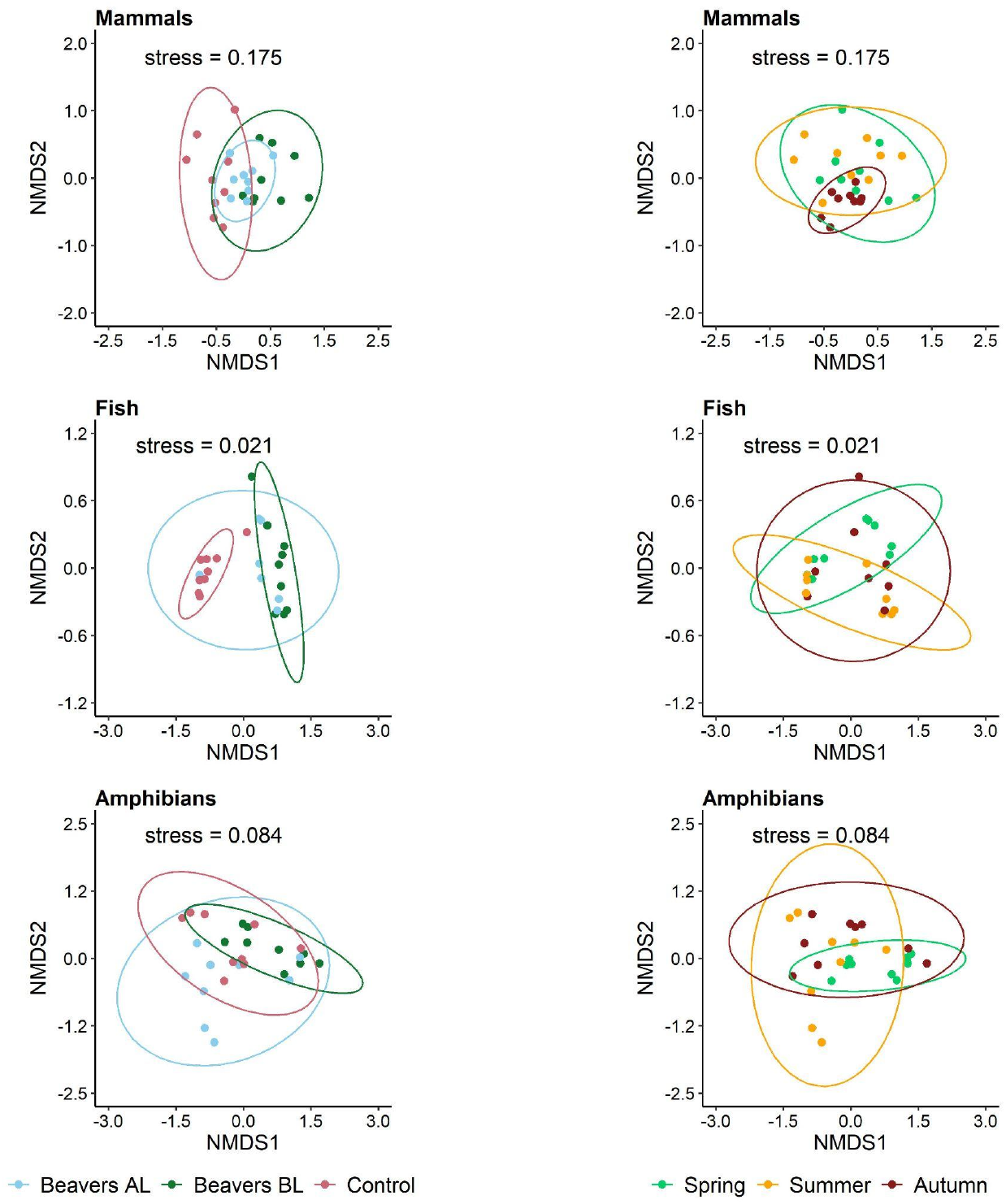
Numerical multi-dimensional scaling of individual sites across all locations and seasons outlining the overlap of multi-dimensional space occupied by the three different sampling groups the analysis is based on Bray-Curtis dissimilarity for fish and amphibians (taking into account variation in relative read counts) and in Jaccard index (based on presence/absence of species) for mammals

### Invertebrate diversity

A total of 2.05m, 1.48m, and 1.92m DNA sequences could be assigned to invertebrate taxa in spring, summer and autumn respectively. Across the data set 486 taxa were identified largely to the species level (n=435, 90%) with some genus level assignments and higher-level assignments (n=51, 10%). The latter were omitted from further analysis. Read counts for species assigned to target taxa in control samples, including field blanks, extraction blanks, PCR negatives, and PCR positives, were generally low. Across 33 control samples, 13 species were detected, each with fewer than 20 reads per sample. Only the two most common species, *Baetis rhodani* and *Eudasyphora cyanicolor*, showed higher read counts (up to 35; Figure S2). These two species were therefore removed from further analyses, and records with fewer than 20 reads per sample were also excluded. After filtering, 365 taxa were retained in the dataset, representing 21 orders across seven higher taxonomic levels (Fig. 6).

**Figure 6:**
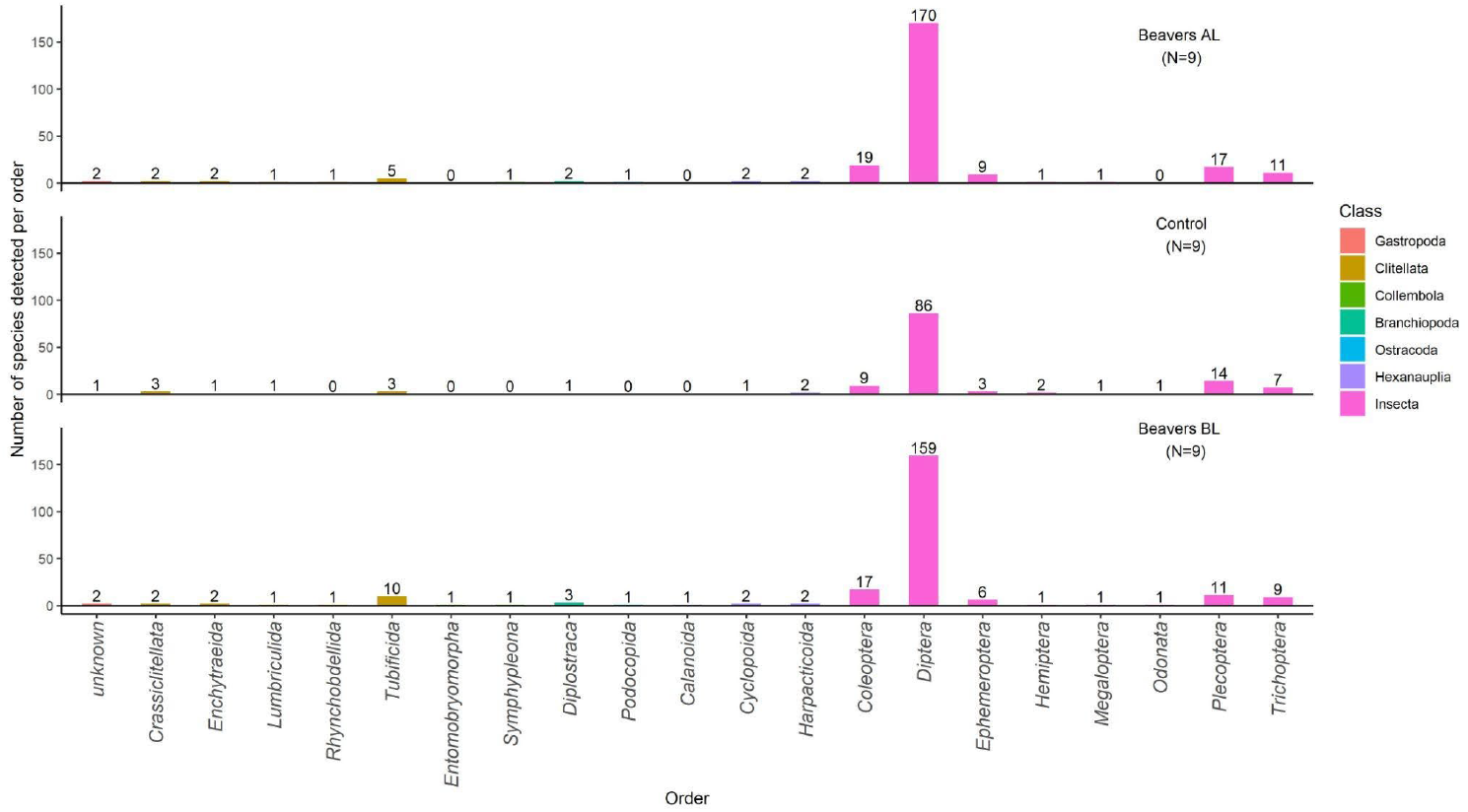
Number of invertebrate species detected for different families and orders in the three spatial groups.Beaver dammed sites above loch (top), control site above loch (middle) and beaver dammed sites below loch (bottom).

The total number of taxa differed among spatial groups with highest gamma diversity in the two beaver impacted groups (249 and 234 species respectively) compared to 136 species detected in the control site.

Mean alpha diversity of samples was significantly different between the three spatial groups (Fig 7; Kruskal-Wallis: X2 = 9.422, DF = 2, P = 0.009), driven by differences between the control and beaver sites. We also observed significant differences between seasons, with lowest species richness being recorded in summer samples (Fig 7; Kruskal-Wallis: X2 = 6.065, DF = 2, P = 0.048).

**Figure 7:**
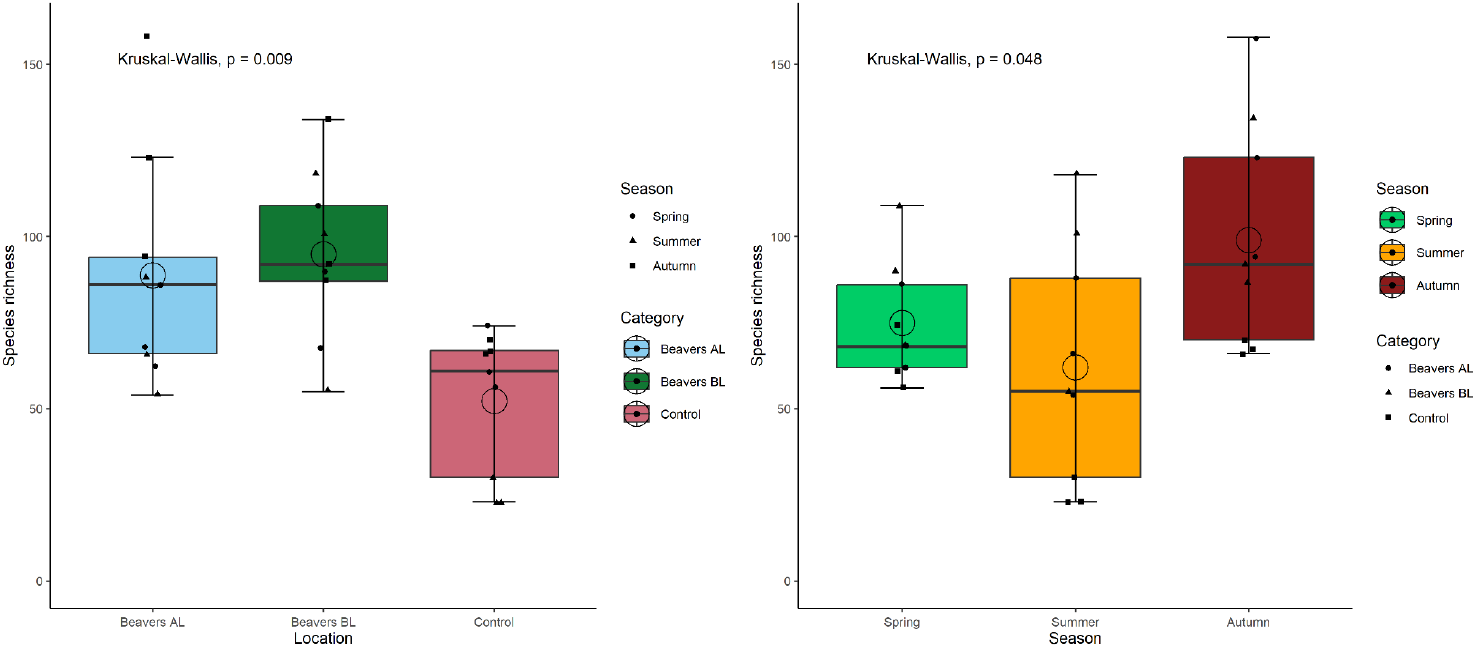
Box plots comparing invertebrate species richness in samples among different spatial groups (left) and different sampling seasons (right).

The invertebrate community composition varied between location and season (Fig. 8). The differences were significant, however the effect of location was stronger (PERMANOVA, R2=0.29, P= 0.001) compared to season (R2=0.20, P= 0.001; Table 2). Multivariate dispersions were homogeneous between locations (ANOVA, P=0.86) however varied between seasons (ANOVA, P= 0.001; Table 2).

**Table 1:** Analysis of invertebrate community similarity among spatial groups and seasons.

|  | Homogeneity of multivariate dispersions (ANOVA) |  |  | Community similarity (PERMANOVA) |  |  |  |
| --- | --- | --- | --- | --- | --- | --- | --- |
| Covariate | df | F | P | df | R2 | F | P |
| Invertebrates |  |  |  |  |  |  |  |
| Location | 2 | 0.14 | 0.864 | 2 | 0.29 | 7.57 | 0.001 *** |
| Season | 2 | 11.6 | <0.001*** | 2 | 0.20 | 5.19 | 0.001 *** |
| Location * Season | n/a | n/a | n/a | 4 | 0.17 | 2.27 | 0.002 ** |

**Figure 8:**
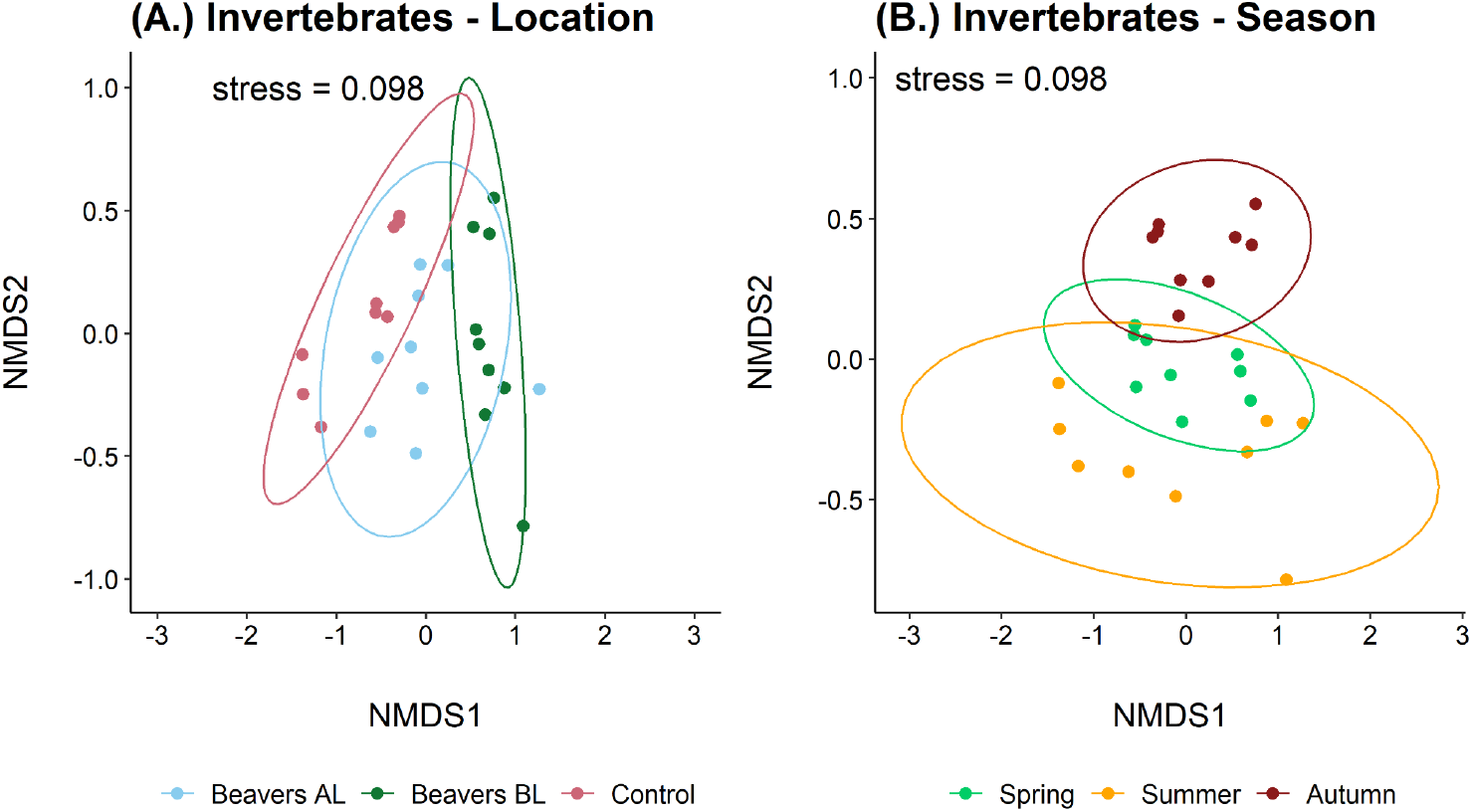
Numerical multi-dimensional scaling of all individual sites across the campaign, outlining the overlap of multi-dimensional space occupied by the three different sampling locations (a.) and seasons (b.) The analysis is based on Jaccards index (based on presence/absence of species) for invertebrate species.

## Discussion

Our results demonstrate that beaver-modified streams supported distinct and, in several respects, more diverse aquatic communities compared to the control stream. Although vertebrate alpha diversity did not differ significantly between spatial groups, overall vertebrate occurrences were higher at beaver sites, with notable increases in amphibian taxa, stickleback, and semi-aquatic mammals such as water vole and water shrew. Community composition differed significantly between beaver and control sites across mammals, fish, and amphibians, with the strongest effects observed for fish communities. Invertebrate responses were even more pronounced, with substantially higher gamma and alpha diversity recorded in beaver-influenced habitats and marked shifts in community composition associated with location. Seasonal effects were also evident, particularly with lower richness observed during summer sampling. Together, these findings suggest that beaver-driven habitat modification strongly influences freshwater biodiversity patterns across multiple taxonomic groups.

### Impact of beaver activity on vertebrate diversity patterns

Vertebrate species richness in individual sample sites (alpha-diversity) and total number of species identified within an experimental group (gamma-diversity) did not differ among beaver impacted locations and control sites without beavers. Furthermore, with a few exceptions the same set of species was identified in all three groups. This was not unexpected given the close proximity of all sample sites and the size and mobility of most of the species recorded here. Most terrestrial vertebrates are likely to roam across the entire enclosure and are likely to be in contact with different waterbodies. Therefore, their presence or absence in individual samples is not necessarily indicative of a specific habitat association. However there is increasing evidence of a range of mammal species using beaver lodge and burrow systems as shelters (Rosell et al., 2025), although at the study site the known structures were in the vicinity of the main loch, and their potential use by other species would probably require more targeted assessments. Fish and amphibians show a low diversity in Scottish upland streams and the species which were found here have broad ecological niches and can utilise a wide range of freshwater habitats. In contrast, studies in more diverse systems have shown that the presence of beavers has a positive effect on the number of fish species (Kemp et al., 2012). Therefore, the lack of difference in the presence of individual species and species richness among spatial groups shown in this study might be due to the overall low species diversity of fish and amphibians in Scottish upland streams and the spatial scale investigated here.

Nevertheless, there were differences in abundance of individual species among different spatial groups as measured by site occupancy and read count. Community similarity analysis showed that the presence of beaver dams has a significant effect on the spatial distribution of fish, amphibian and mammal diversity. The abundance of eDNA from the two main fish species, brown trout and stickleback differed marked between the beaver and control sites. The differences were especially pronounced when comparing the “control stream” with the “beavers below loch” group, which were dominated by just one species each, trout and stickleback respectively. In the “beavers above loch” group the impact on trout was weaker with a reduction of trout sequence read count by approximately 35% compared to the “control stream”. The study of (Needham et al., 2021) provides electro-fishing data from these two spatial groups and an opportunity to put the eDNA results in the context of the conventional approaches for fish monitoring. However, the electrofishing data from Needham *et al*. (2021) were collected in 2014 and 2015, hence the comparability is limited. They found that trout density differed in only one of the two survey years between the two groups with density being higher in the control stream in 2015. While this shows that the eDNA and electro-fishing data are broadly compatible they are not directly comparable due to the different information content of the two approaches. eDNA metabarcoding can provide ecologically meaningful semi-quantitative data (Di Muri, Lawson Handley & Bean, 2020; Griffiths et al., 2020; Nakagawa et al., 2022), but it is difficult to infer absolute abundances from them. This approach becomes less useful in species poor communities. In contrast, electro-fishing surveys can show strong biases towards certain species but can provide information on absolute abundance and age structure. For example, Needham *et al*. (2021) could show a significant change in age structure between the “control stream” and “beavers above loch” group, with the latter having a more diverse size class range and more larger trout being present.

Beaver wetlands provide an array of habitats for small mammals and have been shown to have a positive effect at both the species (Puttock et al., 2023) and community level (Sundell, Liao & Nummi, 2021; Wikar, Ciechanowski & Zwolicki, 2023). In the current study water vole (*A. amphibius*) were seen to benefit from beaver modifications with 9/9 and 9/9 detections in the beaver above and beaver below sites respectively compared with 4/9 detections in the control stream. Similarly, Puttock *et al*., (2023) demonstrated positive co-distribution between beavers and water vole following the River Otter beaver trial. They suggested that beaver wetlands provide more edge habitat for burrowing, felled trees would promote the emergence of riparian vegetation for food and finally, the increased heterogeneity could provide refuge from predators (Puttock et al., 2023).

Beaver modifications also benefited amphibians in the current study, common frogs were detected in higher read counts which suggests higher abundances and palmate newts (*Lissotriton helveticus*) had higher occupancy within beaver modified sites with 9/9 detections at both beaver sites compared to only 4/9 in the control stream. Beaver dams increase water retention creating a more stable hydroperiod for amphibian breeding (Romansic et al., 2021) where the higher temperatures, improved invertebrate productivity, predator refuge and habitat heterogeneity increase amphibian survival and growth rates (Stevens, Paszkowski & Scrimgeour, 2006; Dalbeck, Lüscher & Ohlhoff, 2007; Dalbeck, Janssen & Völsgen, 2014). Dalbeck *et al*., (2009) found that the presence of fish had a negative impact on palmate newts in both beaver and artificial ponds however due to their heterogeneity, beaver ponds provided better refuge for newts and their larvae. Therefore, the increased detections of palmate newts in beaver ponds from the current study could be a result of increased predator refuge from brown trout.

### Impact of beaver activity on invertebrate diversity patterns

Invertebrate species richness in individual samples also did not differ among experimental groups. However, in contrast to vertebrates there was a marked difference in total invertebrate species richness among spatial groups with approximately 78% more species detected at beaver sites than control stream sites. Most previous studies examining the response of macroinvertebrate communities to habitat modification by beavers have reported either lower or similar species richness in beaver ponds compared to unmodified stream habitats (see (Law et al., 2016) for discussion). However, these findings may partly reflect the limited taxonomic resolution of traditional morphological surveys, which often identify organisms only to genus or family level. This is particularly relevant for taxa that are common and diverse in lentic habitats, such as Chironomidae and Tubificidae. In contrast, the molecular approach used here provided species-level identification across a broad range of arthropod families, allowing finer-scale differences in community composition to be detected. Furthermore, eDNA based approaches integrate data over a larger spatial area due to the dispersion of eDNA from the organism that shed it. As a result, the detected communities are more likely to reflect the diversity of habitats present across the entire spatial group, including both ponded (lentic) and flowing (lotic) sections within beaver-modified streams. The greater species richness observed at beaver sites therefore likely reflects the increased habitat heterogeneity created by beaver activity, which produces a lentic–lotic habitat mosaic compared to the more uniformly lotic character of streams without beavers (Law et al., 2017). For example, many species of water beetle and midges are standing water specialists and unlikely to be found in running water. Beaver activity may also enhance the quality of downstream lotic habitats for many species, for example through the action of beaver dams filtering and reducing suspended sediments (Geris, Dimitrova-Petrova & Wilkinson, 2020)

There was also a considerable and significant difference in overall community structure between the different spatial groups as evidenced shown by the PERMANOVA tests. This indicates that the change in habitat structure caused by beaver dams also adds a significant element of beta-diversity on a larger scale across the entire site. This is consistent with other studies including the two case studies from Scotland that reported significant differences in invertebrate communities between unmodified streams and beaver impacted habitats (Law et al., 2016; Needham et al., 2021). Again, the study of Needham *et al*. (2021) provides the most relevant comparison as it was carried out in the same sites. Indeed, the effect of beaver modification on invertebrate community structure reported in our study (*R*^2^=0.29) was very similar to that from Needham *et al*. (2021) (*R*^2^=0.26). This indicates that the eDNA metabarcoding approach can effectively identify changes in macroinvertebrate communities relating to beaver induced habitat modification and is therefore potentially suitable for large-scale monitoring programmes.

### Seasonal variation in eDNA based biodiversity measures

Both the vertebrate and invertebrate data sets showed seasonal variation in biodiversity patterns. The number of species recorded per sample was lowest in summer and the summer samples showed a significantly larger within-group multivariate dispersion i.e. there was greater community composition dissimilarity among summer samples than among spring and autumn samples. Both of these patterns indicate that vertebrate and invertebrate eDNA has a more localised distribution in summer compared to spring and autumn. This has previously been reported for fish eDNA in lakes and rivers (Lawson Handley et al., 2019; Suzuki, Nakano & Kobayashi, 2022). There are several mechanisms that might underlie such seasonal variation. Studies on individual species have demonstrated eDNA shedding rates can vary considerably across seasons due to variation in metabolic rates e.g. mussels (Wacker et al., 2019) or life cycle dynamics, e.g. amphibians (Buxton et al., 2017). The temperature dependency of DNA degradation also adds to the seasonal variation in eDNA detection patterns (Strickler, Fremier & Goldberg, 2015). DNA dispersion is linked to hydrological dynamics which in turn show seasonal variation, for example stratification in lakes or river flow levels. Given the wide taxonomic and ecological range of taxa surveyed in this study it is less likely that the results were biased by the biology of individual species. The lower diversity in summer samples and the stronger differentiation between individual samples is probably due to the increased temperature and low flow rates. This is consistent with a recent study from this system which integrated a published eDNA transport model (Pont, 2024) to reveal the influence of seasonal environmental variation on beaver eDNA transport distances (Macarthur et al., 2026). Irrespective of the causes underpinning these patterns, the results have important implications for the application of eDNA based monitoring. A meaningful pattern of temporal change in biodiversity over multiple years can only be obtained, when monitoring results from the same seasons are compared.

## Conclusions

This study demonstrates that eDNA metabarcoding is an effective tool for detecting biodiversity responses to beaver-driven habitat modification in headwater freshwater systems even at a small geographical scale. Beaver activity altered community composition across multiple taxonomic groups and substantially increased macroinvertebrate gamma diversity, supporting the role of beavers as ecosystem engineers that enhance habitat heterogeneity and beta diversity. Although vertebrate species richness remained similar between beaver and control sites, clear shifts in fish, amphibian, and semi-aquatic mammal assemblages were evident, reflecting the ecological influence of lentic-lotic habitat mosaics created by dam building. Seasonal differences in eDNA patterns further emphasise the importance of temporally standardised monitoring. Overall, our findings highlight the value of combining eDNA metabarcoding with restoration ecology to support scalable biodiversity monitoring and inform future beaver management and reintroduction strategies.

## Supporting information

PDF document containing all supporting information for the publication

## Acknowledgements

We are grateful to the landowners, for allowing site access and for their full support of the project. Matthew Curran, Vicky Johnson and Sam Beck assisted during sample collection. The work was funded by NatureScot and UHI Inverness.

## Author Contributions

Conceptualization: BH. Developing methods - study design: BH, MG, AT, VP. Conducting research - fieldwork: BH, JM. Conducting research - laboratory work: JM, BM, DS. Data analysis, preparation of figures and tables: NG, JM. Writing – original draft: BH. Writing – review & editing: BH, BM, DS, NG, JM, MG, AT, VP

## Funding

The work was supported by NatureScot and UHI Inverness.

## Conflicts of interest

The authors declare no conflicts of interest

## Data Availability statement

All scripts and corresponding data will be made publicly available at zenodo following the acceptance of this publication

## Notes

### Competing Interest Statement

The authors have declared no competing interest.

