## Supplementary material for "Biodiversity effects of beaver activity in a semi-natural enclosure revealed by eDNA": PDF document containing all supporting information for the publication

### Annexes

#### Annex 1 – Detailed methods

##### Sample collection

2 L water samples each, comprising 5 x 400 ml subsamples taken at 5 m intervals, were collected from the site as described in Hänfling et al. (2016). Samples were collected from the shoreline at equidistant locations (access permitting) using sterile Gosselin™ HDPE plastic bottles (Fisher Scientific UK Ltd, UK) and disposable nitrile gloves (STARLAB, UK). A blank (2 L molecular grade water [MGW]) was transported alongside samples in insulated coolboxes with ice packs. Coolboxes and ice packs were sterilised with 10% v/v chlorine-based commercial bleach (Elliott Hygiene Ltd, UK) solution (containing ~3% sodium hypochlorite) before use.

##### eDNA capture and extraction

Water samples were vacuum-filtered within 24 hrs of collection in a dedicated eDNA facility at UHI Inverness. Surfaces and equipment were sterilised before, during, and after set-up in all work areas. Surfaces and vacuum pumps were wiped with 10% bleach solution. Non-electrical equipment was immersed in 10% bleach solution for 10 minutes, followed by 5% v/v MicroSol detergent (Anachem), and rinsed with purified water. Each 2-l water sample was filtered through sterile 0.45 µm mixed cellulose ester membrane filters with pads (47 mm diameter; Whatman, GE Healthcare) using Pall Laboratory Manifold (516-1038) filtration units. Two filters were used for each sample and 30 mins allowed for water to pass through each filter, totalling one hour of filtration time per sample. Blanks (n = 9) transported alongside samples were filtered during the last round of filtration for each day. Equipment was sterilised after each round of filtration. Filters were removed from pads using sterile tweezers and placed in sterile 5 ml polypropylene screw-cap tubes (Axygen, Fisher Scientific UK Ltd.), placed in gripseal bags, and stored at -20°C until DNA extraction. DNA extraction followed the Mu-DNA water protocol (Sellers *et al.*, 2018). Briefly, captured DNA was liberated from the filter membranes via bead-milling lysis. The lysate underwent an inhibitor removal step prior to purification with silica membrane EZ-10 DNA Mini Spin Columns (NBS Biologicals) and final elution (100 µl). An extraction blank (n = 8), consisting only of extraction buffers and sterile garnet beads, was extracted alongside samples. 2 µl aliquots of each DNA extract were taken for measurement with a Qiaxpert spectrophotometer (Qiagen) to assess yield and purity. 20ul working aliquots were taken from the stock tube this ensured stock tubes did not undergo continual freeze thawing. DNA extracts were then frozen at -20°C until PCR amplification.

##### Vertebrate DNA metabarcoding library preparation

Dedicated rooms were available for pre-PCR and post-PCR processes. Pre-PCR processes were performed in the UHI Inverness eDNA facility, which has separate rooms for filtration, DNA extraction, and PCR preparation of sensitive environmental samples. PCR reactions were set up in an ultraviolet (UV) and bleach sterilised laminar flow hood. Post-PCR processes were performed in the UHI Inverness IBFC laboratory, which has rooms dedicated to pre-PCR of nonsensitive samples, PCR, agarose gel electrophoresis, PCR product purification, library quality control (Qubit, Agilent Zero Agarose Gel (ZAG) DNA Analyzer system, real-time quantitative PCR (qPCR)), and sequencing (Illumina MiSeq).

Libraries were prepared for sequencing using a nested metabarcoding workflow with a two-step PCR protocol, where Multiplex Identification (MID) tags (unique 8-nucleotide sequences) were included in the first and second PCR for sample identification (Kitson et al. 2019). DNA extracts were PCR-amplified using vertebrate-specific primers that target a 106 bp fragment of the mitochondrial 12S ribosomal RNA (rRNA) region in fish (Riaz et al. 2011). The primers were modified for the present study to include MID tags, heterogeneity spacers, sequencing primers, and pre-adapters. There were 24 unique MID tags for the forward and 24 unique MID tags for the reverse primers. This allowed 24 samples to each be labelled with a unique forward and a unique reverse primer to reduce barcode misassignment and tag jumps (Deakin *et al.*, 2014; Schnell, Bohmann & Gilbert, 2015). During the first PCR, samples were processed in batches (i.e. up to 20 eDNA samples, one filtration blank, one extraction blank, one negative control and one positive control). The PCR positive control was zebra mbuna (*Maylandia zebra*) DNA (0.05 ng/µl). M. zebra is an exotic cichlid which is not found in UK freshwater habitats.

The first PCR was performed in triplicate for each sample/control to combat stochasticity arising from low target DNA concentrations. PCR replicates for each sample/control had the same tag combination. PCR reactions were performed in 15 µl volumes, consisting of: 7.5 µl of Q5® High-Fidelity 2x Master Mix (New England Biolabs), 0.3 µl of Thermo Scientific Bovine Serum Albumin (Fisher Scientific UK Ltd.), 3.7 µl of MGW (Fisher Scientific UK Ltd.), 1.8 µl of each 10 µM tagged primer, and 1.7 µl of template DNA. PCR reactions were sealed with strip caps (Fisher Scientific). PCR was performed on an Agilent Surecycler 8800 with the following thermocycling profile: 98°C for 5 mins, 35 cycles of 98°C for 10 s, 58°C for 20 s and 72°C for 30 s, 72°C for 7 mins then held at 4°C.

PCR products were stored at 4°C until PCR technical replicates for each sample/control were pooled. 2 µl of each pooled PCR product was visualised using an Agilent ZAG DNA analyzer. PCR products were deemed positive where there was amplification at the expected size (200-300 bp) on the gel. PCR products were stored at -20°C until they were normalised and pooled to create sub-libraries for a double-size selection bead purification protocol. Sub-libraries were then purified using Pro-Nex size selection beads (Promega) using 75 µL of magnetic beads to a 50 µL DNA sample. Eluted DNA (25 µl) was stored at 4°C until second PCR amplification.

The second PCR bound pre-adapters, MID tags, and Illumina adapters to the purified sub-libraries. 10 unique forward and reverse MID tag combinations were selected and applied to 39 sub-libraries split across four sequencing runs. Two replicates were performed for each sub-library in 50 µl volumes, consisting of: 25 µl of Q5 High-Fidelity 2x Master Mix (New England Biolabs), 13 µl of MGW (Fisher Scientific UK Ltd.), 3 µl of each 10 µM tagged primer (final concentration 0.6 µM; Integrated DNA Technologies), and 4 µl of template DNA. PCR was performed on an Agilent Surecycler 8800 with the following thermocycling profile: 95°C for 3 mins, 10 cycles of 98°C for 20 s and 72°C for 1 min, 72°C for 5 mins then held at 4°C. PCR duplicates for each sub-library had the same tag combination.

PCR products were stored at 4°C until duplicates for each sub-library were pooled. 2 µl of each pooled PCR product was visualised on an Agilent ZAG DNA analyzer. PCR products were deemed positive where there was amplification at the expected size (300-400 bp) on the gel. Sub-libraries were stored at 4°C until double-size selection bead purification. A second purification was then carried out on the PCR2 products with Pro-Nex size selection beads (Promega) using 60 µL of beads to a 50 µL DNA sample. Eluted DNA (25 µl) was stored at 4°C until normalisation and final purification.

Sub-libraries were quantified on a Qubit 3.0 fluorometer using a dsDNA HS Assay Kit (Invitrogen) and normalised by pooling according to sample size and library concentration. The pooled library was purified using the same ratios, volumes, and protocol as the second PCR purification. Based on the Qubit™ concentration, the library was diluted to 4 nM. The library was then quantified by qPCR using the KAPA Library Quantification Kit for Illumina

(Roche). Based on the qPCR concentration, the library was adjusted to 4 nM and denatured following the Illumina MiSeq library denaturation and dilution guide. The final library was sequenced at 13 pM with 10% PhiX Control on an Illumina MiSeq using 2 x 300 bp V3 chemistry (Illumina).

##### Vertebrate bioinformatics

Sequencing data was automatically demultiplexed to separate (forward and reverse) fastq files per library using the onboard Illumina MiSeq Reporter software. Library sequence reads were further demultiplexed to sample using a custom Python script. *Tapirs*, a reproducible workflow for the analysis of DNA metabarcoding data (https://github.com/EvoHull/Tapirs), was used for taxonomic assignment of demultiplexed sequencing reads. *Tapirs* uses the *Snakemake* workflow manager (Köster & Rahmann, 2012) and a *conda* virtual environment to ensure software compatibility.

Raw reads were quality trimmed from the tail with a 5 bp sliding window (qualifying phred score of Q30 and an average window phred score of Q30) using *fastp* (Chen *et al.*, 2018), allowing no more than 40% of the final trimmed read bases to be below Q30. Primers were removed by trimming the first 18 bp of both forward and reverse reads. Reads were then tail cropped to a maximum length of 106 bp and reads shorter than 90 bp were discarded.

Sequence read pairs were merged into single reads using *fastp*, provided there was a minimum overlap of 20 bp, no more than 5% mismatches and no more than 5 mismatched bases between pairs. Only forward reads were kept from read pairs that failed to be merged. A final length filter removed any reads longer than 110 bp to ensure sequence lengths approximated the expected fragment size (~106 bp).

Redundant sequences were removed by clustering at 100% read identity and length (--derep_fulllength) in *VSEARCH* (Rognes *et al.*, 2016). Clusters represented by less than three sequences were omitted from further processing. Reads were further clustered (--cluster_unoise) to remove redundancies due to sequencing errors (retaining all cluster sizes). Retained sequences were screened for chimeric sequences with *VSEARCH* (--uchime3_denovo).

The final clustered, non-redundant query sequences were then compared against a curated UK vertebrate reference database (Harper *et al.*, 2018) using BLAST (Zhang *et al.*, 2000). Taxonomic identity was assigned using a custom majority lowest common ancestor (MLCA) approach based on the top 2% query BLAST hit bit-scores, with at least 90% query coverage and a minimum identity of 98%. Of these filtered hits, 80% of unique taxonomic lineages therein had to agree at descending taxonomic rank (domain, phylum, class, order, family, genus, species) for it to be assigned a taxonomic identity. If a query had a single BLAST hit it was assigned directly to this taxon only if it met all MLCA criteria. Read counts assigned to each taxonomic identity were calculated from query cluster sizes. Lowest taxonomic rank was to species and assignments higher than order were classed as unassigned. Following taxonomic assignment, a minimum threshold of five reads was applied to remove low-frequency reads alongside a species-specific contaminant threshold to remove any reads in samples which were lower than detections in controls (Figure S1) (Macarthur *et al.*, 2025). Most reads were assigned to the species level, but as the molecular marker used here cannot distinguish certain species reliably, the reads belonging to these species were assigned to the next possible higher taxonomic level. Reads assigned to positive controls, reads which could not be assigned to any taxon and samples with no taxonomically assignable reads were also removed from the data set.

##### Invertebrate eDNA metabarcoding library preparation

Dedicated rooms were available for pre-PCR and post-PCR processes. Pre-PCR processes were performed in the UHI Inverness eDNA facility, which has separate rooms for filtration, DNA extraction, and PCR preparation of sensitive environmental samples. PCR reactions were set up in an ultraviolet (UV) and bleach sterilised laminar flow hood. Post-PCR processes were performed in the UHI Inverness IBFC laboratory, which has rooms dedicated to pre-PCR of nonsensitive samples, PCR, agarose gel electrophoresis, PCR product purification, library quality control (Qubit, Agilent Zero Agarose Gel (ZAG) DNA Analyzer system, real-time quantitative PCR (qPCR)), and sequencing (Illumina MiSeq).

Libraries were prepared for sequencing using a two-step PCR protocol. The first PCR amplified the target region with Multiplex Identification (MID) tags (unique 8-nucleotide sequences) were included in the second PCR for sample identification. DNA extracts were PCR-amplified using primers that target a ~140 bp fragment of the cytochrome c oxidase subunit 1 (COI) mitochondrial region in invertebrates (Leese *et al.*, 2021). The primers were modified for the present study to include heterogeneity spacers, sequencing primers, and pre-adapters. During all stages, samples were processed in the same batches of up to 22 eDNA samples, one negative control and one positive control, keeping sampling replicates in the same batch where possible. The PCR positive control was bluebottle fly (*Calliphora vicina*) genomic DNA (0.05 ng/µl).

The first PCR was performed in triplicate for each sample/control to combat stochasticity arising from low target DNA concentrations. PCR reactions were performed in 25 µl volumes, consisting of: 12.5 µl of Multiplex PCR Master Mix (QIAGEN Multiplex PCR Plus Kit), 0.5 µl of Thermo Scientific Bovine Serum Albumin (Fisher Scientific UK Ltd.), 7 µl of MGW (Fisher Scientific UK Ltd.), 1.5 µl of each 10 µM tagged primer, and 2 µl of template DNA. PCR reactions were sealed with strip caps (Fisher Scientific). PCR was performed on an Agilent Surecycler 8800 with the following thermocycling profile: 95°C for 5 mins, 35 cycles of 95°C for 30 s, 50°C for 90 s, 72°C for 2 mins, and a final elongation of 68°C for 10 mins then held at 4°C.

PCR products were stored at 4°C until PCR technical replicates for each sample/control were pooled. 5 µl of each pooled PCR product was visualised on an Agilent ZAG DNA analyzer. Of all the samples/controls processed, only the positive control was visibly amplified at the expected size (~260 bp) after PCR. All PCR products were individually purified with a size selection bead purification protocol (Rohland & Reich, 2012). A second purification was then carried out on the PCR2 products with Pro-Nex size selection beads (Promega) using 60 µL of beads to a 50 µL DNA sample. Eluted DNA (25 µl) was stored at 4°C until second PCR amplification.

The second PCR bound pre-adapters, MID tags, and Illumina adapters to the purified PCR products. Each sample/control was treated with one of the 384 unique forward and reverse MID tag combinations (16 forward and 24 reverse). Two replicates were performed for each sample/control in 50 µl volumes, consisting of: 25 µl of Q5 High-Fidelity 2x Master Mix (New England Biolabs), 13 µl of MGW (Fisher Scientific UK Ltd.), 3 µl of each 10 µM tagged primer (final concentration 0.6 µM; Integrated DNA Technologies), and 4 µl of template DNA. PCR was performed on an Applied Biosystems Veriti Thermal Cycler (Life Technologies) with the following thermocycling profile: 95°C for 3 mins, 10 cycles of 98°C for 20 s, 72°C for 1 min, and a final elongation of 72°C for 5 mins then held at 4°C. PCR duplicates for each sample/control had the same tag combination.

PCR products were stored at 4°C until PCR technical replicates for each sample/control were pooled. 5 µl of each pooled PCR product was visualised on an Agilent ZAG DNA analyzer. PCR products were deemed positive where there was amplification at the expected size (~330 bp) on the gel. Each processed batch of PCR products were pooled according to band strength (no/very faint band = 20 µl, faint band = 15 µl, bright band = 10 µl, very bright band = 5 µl) on gel (Alberdi *et al.*, 2018) to create sub-libraries for a size selection bead purification protocol (Rohland & Reich, 2012). A second purification was then carried out on the PCR2 products with Pro-Nex size selection beads (Promega) using 60 µL of beads to a 50 µL DNA sample. Eluted DNA (25 µl) was stored at 4°C until normalisation and final purification.

Sub-libraries were quantified on a Qubit 3.0 fluorometer using a dsDNA HS Assay Kit (Invitrogen) and normalised by pooling according to sample size and library concentration. Based on the Qubit™ concentration, the library was diluted to 4 nM. The library was checked with an Agilent ZAG DNA Analyzer to verify a fragment of the expected size (~330 bp) remained. The library was then quantified by qPCR using the KAPA Library Quantification Kit for Illumina (Roche). Based on the qPCR concentration, the library was adjusted to 4 nM and denatured following the Illumina MiSeq library denaturation and dilution guide. The final library was sequenced at 13 pM with 10% PhiX Control on an Illumina MiSeq using 2 x 300 bp V3 chemistry (Illumina).

**UK invertebrate reference database creation**

The reference database was created from the curated reference sequences used in Harper et al. (2021) (https://doi.org/10.5281/zenodo.3993125). Harper et al. (2021) did not include dipterans in their database due to GenBank records for Diptera missing important record features. After initial tests we discovered errors in assignments, the PCR positive (a fly) was assigned to a Coleoptera species. To remedy these issues, additional records for UK Diptera species were downloaded from The Barcode of Life Data System (BOLD) (https://www.boldsystems.org/index.php/). BOLD records are more tightly curated than Genbank, and are focused on a few specific regions, in particular, COI. At present, the *Tapirs* workflow uses NCBI taxonomy, so only BOLD reference sequences with an associated NCBI accession number were used for database creation. An NCBI taxonomic id (taxid) was acquired for each reference sequence to create an accession-to-taxid map. The final *BLAST* database was created from the reference sequences using *makeblastdb* (see https://www.ncbi.nlm.nih.gov/books/NBK569841/) and included the accession-to-taxid map (-*taxid_map*) to allow for taxonomic assignment of OTUs for downstream *Tapirs* analysis.

**Invertebrate eDNA metabarcoding bioinformatics**

Sequencing data was automatically demultiplexed to separate (forward and reverse) fastq files per sample using the onboard Illumina MiSeq Reporter software. *Tapirs*, a reproducible workflow for the analysis of DNA metabarcoding data (https://github.com/EvoHull/Tapirs), was used for taxonomic assignment of demultiplexed sequencing reads. *Tapirs* uses the *Snakemake* workflow manager (Köster & Rahmann, 2012) and a *conda* virtual environment to ensure software compatibility.

Raw reads were quality trimmed from the tail with a 5 bp sliding window (qualifying phred score of Q30 and an average window phred score of Q30) using *fastp* (Chen *et al.*, 2018), allowing no more than 40% of the final trimmed read bases to be below Q30. Primers were removed by trimming the first 26 and 23 bp of forward and reverse reads respectively. Reads were then tail cropped to a maximum length of 142 bp and reads shorter than 100 bp were discarded.

Sequence read pairs were merged into single reads using *fastp*, provided there was a minimum overlap of 20 bp, no more than 5% mismatches and no more than 5 mismatched bases between pairs. Only forward reads were kept from read pairs that failed to be merged. A final length filter removed any reads longer than 160 bp to ensure sequence lengths approximated the expected fragment size (~142 bp).

Redundant sequences were removed by clustering at 100% read identity and length (--*derep_fulllength*) in *VSEARCH* (Rognes *et al.*, 2016). Clusters represented by less than three sequences were omitted from further processing. Reads were further clustered (--*cluster_unoise*) to remove redundancies due to sequencing errors (retaining all cluster sizes). Retained sequences were screened for chimeric sequences with *VSEARCH* (--*uchime3_denovo*).

The final clustered, non-redundant query sequences were then compared against the UK invertebrate reference database using *BLAST (Zhang et al., 2000)*. Taxonomic identity was assigned using a custom majority lowest common ancestor (MLCA) approach based on the top 2% query *BLAST* hit bit-scores, with at least 90% query coverage and a minimum identity of 95%. Of these filtered hits, 80% of unique taxonomic lineages therein had to agree at descending taxonomic rank (domain, phylum, class, order, family, genus, species) for it to be assigned a taxonomic identity. If a query had a single *BLAST* hit it was assigned directly to this taxon only if it met all previous MLCA criteria. Read counts assigned to each taxonomic identity were calculated from query cluster sizes. Lowest taxonomic rank was to species and assignments higher than order were classed as unassigned.

### Annex 2 - additional analyses

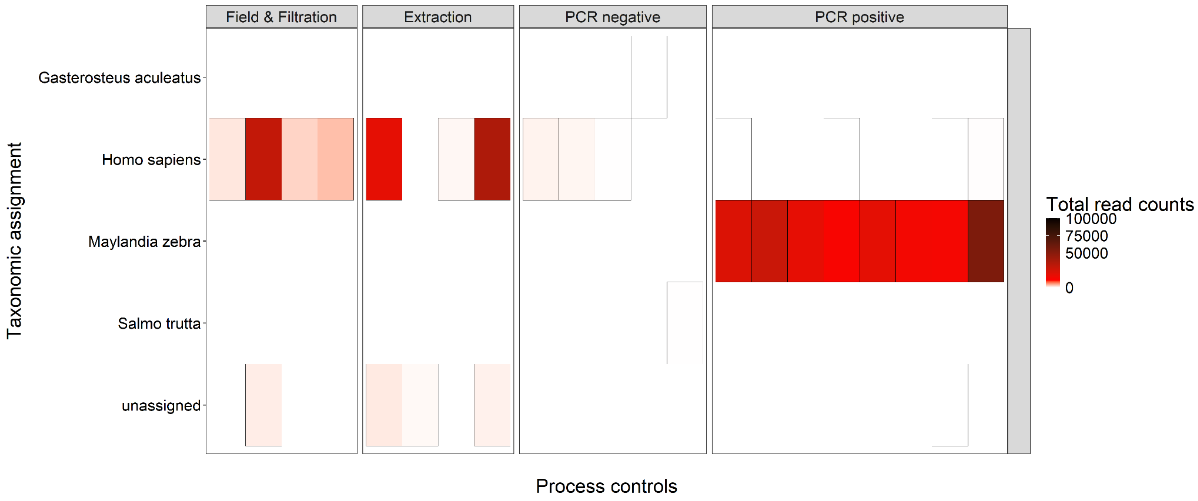


Figure S1: Heatmap of total read counts across control samples collected during different lab and field processes from the vertebrate assay.

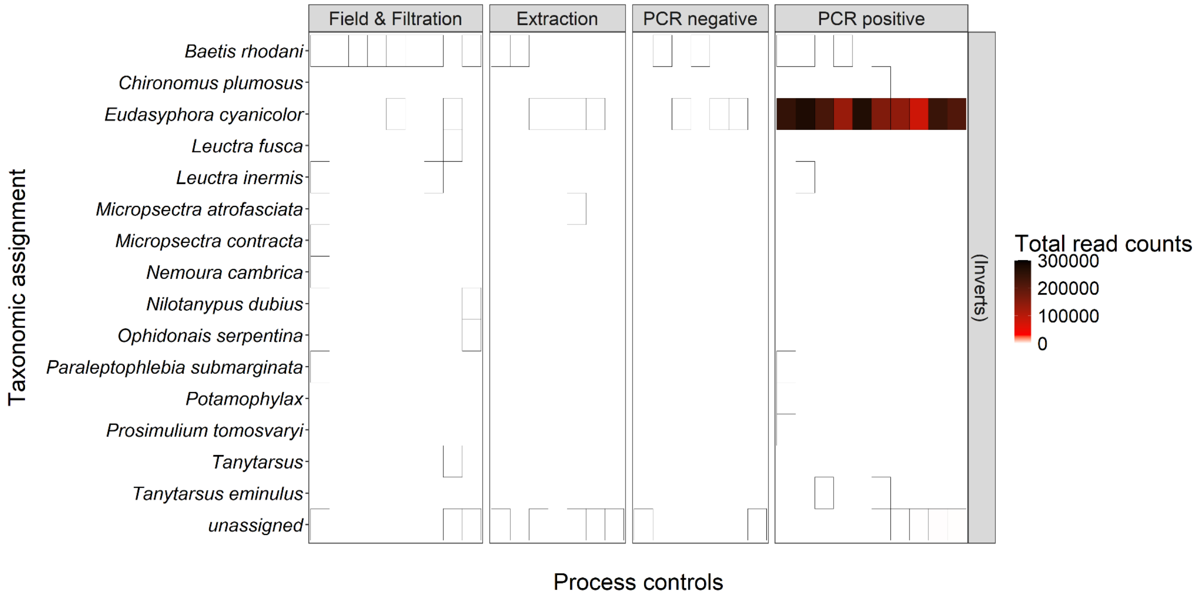


Figure S2: Heatmap of total read counts across control samples collected during different lab and field processes from the invertebrate assay.
